# Inflect: Optimizing Computational Workflows for Thermal Proteome Profiling Data Analysis

**DOI:** 10.1101/2020.10.31.363523

**Authors:** Neil A. McCracken, Sarah A. Peck Justice, Aruna B. Wijeratne, Amber L. Mosley

## Abstract

The use of CETSA and Thermal Proteome Profiling (TPP) analytical methods are invaluable for the study of protein-ligand interactions and protein stability in a cellular context. These tools have increasingly been leveraged in work ranging from understanding signaling paradigms to drug discovery. Consequently, there is an important need to optimize the data analysis pipeline that is used to calculate protein melt temperatures (T_m_) and relative melt shifts from proteomics abundance data. Here we report a user-friendly analysis of the melt shift calculation workflow where we describe the impact of each individual calculation step on the final output list of stabilized and destabilized proteins. This report also includes a description of how key steps in the analysis workflow quantitatively impacts the list of stabilized/destabilized proteins from an experiment. We applied our findings to develop a more optimized analysis workflow that illustrates the dramatic sensitivity of chosen calculation steps on the final list of reported proteins of interest in a study and will make the R based program Inflect available for research community use. Overall, this work provides an essential resource for scientists as they analyze data from TPP and CETSA experiments and implement their own analysis pipelines geared towards specific applications.

## INTRODUCTION

Within the complex cellular milieu, there has long been an inability to screen for untargeted changes in protein stabilization or de-stabilization. The advent of Cellular Thermal Shift Analysis (CETSA) ^1^ and thermal proteome profiling (TPP) ^2, 3^ has rapidly increased our ability to measure changes in protein stability with the context of the intact proteome. The CETSA / TPP workflow begins with culture of cells exposed to different conditions such as treated with a small molecule vs. vehicle or that have different genetic backgrounds.^4, 5^ After culture, the cells are lysed in a non-denaturing extraction buffer and the cellular debris is pelleted and discarded. Supernatant is decanted, aliquoted, and subsequently exposed to a thermal gradient (typically using a PCR machine) ranging from ambient to 90°C. Alternatively, intact cells can be exposed to a thermal gradient.^3^ During this heat treatment, the bulk of the proteins in the solution unfold at a temperature range based on their inherent biophysical properties such as individual structure and their interactions with other partners (including proteins, small molecules, metabolites, etc.). As they unfold, proteins have a greater propensity for aggregation with nearby unfolded proteins and may also precipitate post-aggregation. After the short thermal treatment, the heat-treated samples are centrifuged to pellet aggregated protein. The supernatant containing the soluble fraction is decanted once again, proteolytically digested, cleaned up and labeled with isobaric chemical tags such as tandem mass tag (TMT) reagents for multiplexed analysis by LC-MS/MS. Relative peptide fragment abundance values from a LC-MS/MS experiment are analyzed using a proteomics search program and the list of reported protein melt curves is further processed manually or by an analysis program to yield a list of proteins affected in the experiment. Each step in this described computational process, while having an essential role in the execution of the assay, also has its own potential for adding variability to the final output and conclusions from the study. Challenges with accounting for variability were addressed in part by R and Python based pipelines that calculate the melt shift curves from proteomics program dataset ^6, 7^. The pipeline that accompanied the TPP method, hereafter described as “TPP-TR”, uses raw abundance values from a proteomics program and processes the data through several steps prior to calculating melt temperatures (T_m_) and by comparison melt shift values. The operations used in these steps include data filtering, normalization, meltome quantification by curve fitting with correction, individual protein melt fitting, along with melt temperature and shift calculations.

Despite the availability of resources like TPP-TR that do the heavy lifting in melt shift analysis, there has been no report to date that describes how the chosen analysis steps can impact the final study conclusions. Along with this void in analysis, there have also been very recent reported challenges with TPP methods that can be traced to differences in computational analysis ^8-10^. Differences in observations between groups provide evidence that more work is needed to better understand potential gaps in the computational workflow that remain to be optimized for specific TPP applications. In order to address these deficiencies, we investigated the existing TPP-TR workflow with aims to better describe and optimize the output of a TPP experiment. Herein we describe the relative impact of each melt shift analysis step on the total number of proteins and melt shift standard deviations. We also used our findings to develop an analysis workflow that acts as a complementary pipeline to the existing TPP-TR. We are making our R based analysis pipeline, named “Inflect”, available to the community in order that it can be utilized to aid in gaining more complementary results for comparison to results that can already be obtained with other analysis programs. Our findings summarized below will allow researchers to not only better leverage the results from the costly and time consuming TPP experiments, but also act as a resource for those who develop their own algorithms for analysis.

## MATERIALS AND METHODS

### Datasets

*Peck Justice Data Set -* The first data set used in our analysis is one where the investigators illustrated a novel approach for utilizing the TPP-TR workflow to understand the impact of genetic mutations on the melt of the proteome.^5^ Their data set was generated from *S. cerevisiae* strains, with mutations to the ORFs encoding proteasome subunits Pup2 and Rpn5. The first and third data sets (p1 and p3) generated from a wild type (WT) strain and mutant *pup2* and *rpn5* cells were used in our pipeline analysis. Raw abundance values reported from a search in Proteome Discoverer™ were used in our analysis. *Perrin Data Set -* The second data set used in the analysis was reported by Perrin and co-workers using CETSA to identify targets of Panobinostat in organs and blood of rodents and humans, respectively.^11^ The raw data files were analyzed in Proteome Discoverer™ Version 2.4. Files for the rat kidney and liver were obtained from PRIDE Project ID PXD015427 (sample IDs 02290_F1_R1_P0189540B, 02293_F1_R1_P0189550B, 02066_F1_R1_P0177049B and 02065_F1_R2_P0177039B) while files for the human PBMC and whole blood data sets were obtained from PRIDE Project ID PXD015373 (files 02032_F1_R1_P0175529B and 02604_F1_R1_P0204098E). Proteome Discover searches for the rat data set were searched against *Rattus norvegicus* NCBI 062312, using trypsin enzyme setting, Precursor Mass Tolerance at 20 ppm and fragment Mass Tolerance at 0.5 Da. Regression settings for the search used non-linear Regression with coarse parameter tuning. The same search settings were used for the human blood data sets except the *Homo sapien*s (092919) database from Uniprot was used.

Search results from human whole blood raw data sets were used as the “WT” data sets in the pipelines while the peripheral blood mononuclear cell (PBMC) data sets were designated as the “mutant” data sets. Melt shifts (ΔT_m_) were calculated by subtracting the “mutant” melt temperature from the “WT” melt temperature and destabilized proteins were those with positive shifts while stabilized proteins were those with negative shifts. After searching against the *Rattus norvegicus* proteome, the raw abundance values for each protein were also analyzed using both the TPP-TR and our pipeline in R. The kidney data sets were set as the “mutant” strains while the liver data sets were designated as “WT” in order that each of these two organs could be compared against the liver data sets. The analysis workflows were the same as those that were previously described for the Peck-Justice data set.

### JMP analysis

The statistical analysis software JMP^®^ Pro 14 and Pro 15 were used to randomly vary five of the factors used in the custom TPP analysis workflow along with their respective two-level ranges were used to generate a full factorial design of experiments (DoE). A total of 32 experiment conditions were created and each of the 12 data sets discussed in this report were used to evaluate the performance of each combination of steps (288 total experiments). The outputs of the workflow analyses were both the total number of reported significant proteins along with the standard deviation of the observed melt temperatures. These outputs not only allowed for an understanding of the “signal” that came out of each workflow combination but also gave an appreciation of the level of uncertainty from the overall data. Desirable conditions were those where there were high levels of proteins reported with low levels of standard deviation. A definition of significance was used to find melt curves with calculated R^2^ greater than or equal to 0.95.

### R analysis

RStudio version 1.3.1056 was used for analysis code development and execution. R programs were used firstly for the development of the TPP analysis pipeline that we describe. The optimized workflow “Inflect”, was also coded in RStudio. R programs were also coded for the multivariate analyses that were conducted to determine the relative impact of analysis steps on the final TPP pipeline outputs. R programs were also used to generate various diagrams shown in the figures. GraphPad Prism 8 was used for the generation of plots in the figures section.

### Inflect accessibility

The Inflect code is available at https://github.com/namccrac/Inflect.git. The outputs of the program are as follows: The outputs of this program are as follows: Results.xlsx file - lists the calculated melt shifts and related data for each protein regardless of the criteria (Rsquared and standard deviations); SignificantResults.xlsx file - lists the calculated melt shifts and related data for each protein that was considered significant by the criteria above; Curves Folder - this folder contains the melt curves (in pdf format) for each protein regardless of the significance of the curve; Significant Curves Folder - this folder contains the melt curves (in pdf format) for significant proteins only; Normalized Condition and Control result files - these files contain the normalized abundance values for each protein and at each temperature; Waterfall plot - shows the calculated melt shifts across the proteome in the study. The melt shifts are plotted in order of value (from highest to lowest). A PDF version of this plot is created in the Curves folder. The required inputs are described at the link above.

## RESULTS AND DISCUSSION

Protein melt shift calculation of TPP experimental data can be delineated into 10 steps, which are summarized pictorially in Figure 1. Step 1 excludes data that does not meet pre-defined quality control criteria, followed by step 2 that normalizes the abundance values for each protein to the lowest temperature abundance. Step 3 uses statistics to quantify the total protein meltome and the curve fit routine in step 4 uses non-linear equations to describe the meltome shape. Step 5 calculates correction values based on the actual and predicted curve fit values, after which constants are then used to correct normalized abundance values for each protein. Curve fitting again occurs in step 6, but on each individual protein abundance that has been corrected. The computationally laborious step 6 fits non-linear equations to each of the thousands of individual proteins melt curves followed by another exclusion in step 7 to remove proteins that do not meet another set of quality control criterion. The calculation of the melt temperature for each protein occurs in step 8 after which the melt shift is calculated in step 9. The final step 10 involves the summary of the proteins with significant stabilization or de-stabilization based on the shift of all calculated proteins.We utilized two published CETSA/TPP studies ^11, 12^ with a total of 12 separate data sets to define and quantify the relative impact of each data analysis step on the output from an experiment. The Peck-Justice experiments investigated the impact of genetic mutations in proteasome subunits Pup2 and Rpn5 on protein interactions in *S. cerevisiae* on protein interactions. The Perrin experiments focused on use of CETSA to find Panobinostat targets in human blood and rat organs while also using a 10-temperature gradient (without drug) to probe for interactions in crude cell and tissues. These data sets were chosen because they represent more recent executions of the CETSA / TPP workflows, have publicly available raw data and also feature the use of the TMT label sets. The analysis was demarcated into 10 steps with some key parameters discussed in more detail below.

**Figure 1.**
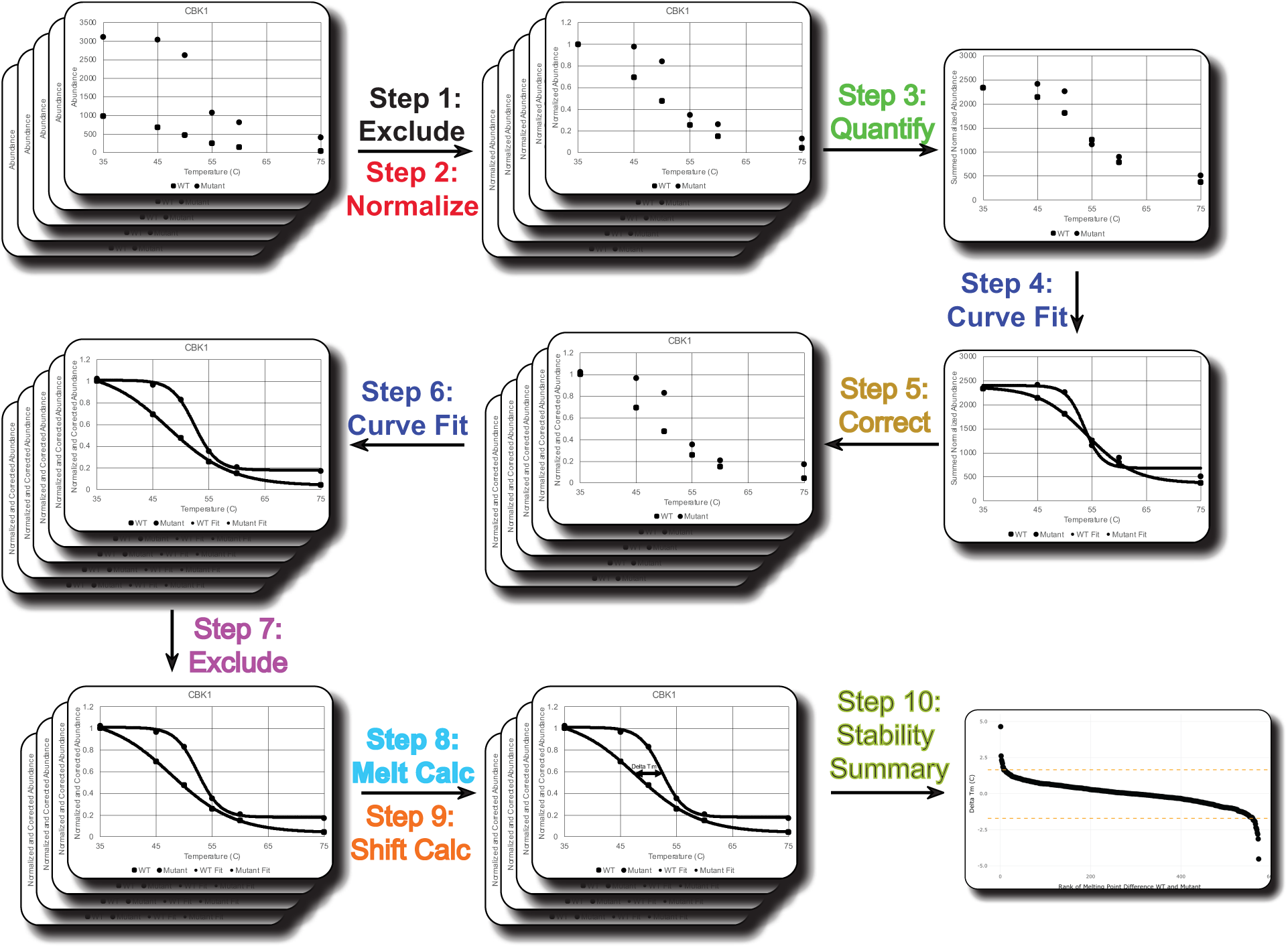
Pictorial representation of the general data analysis workflow from a TPP experiment. Step 1 in the pipeline excludes data while step 2 normalizes the abundance values for each protein. Step 3 quantifies the total protein meltome and the curve fit routine in step 4 uses non-linear equations to describe the meltome shape. Step 5 calculates correction values. Curve fitting occurs again in step 6 while exclusion is used again in step 7 to remove proteins that do not meet fit quality criteria. The calculation of the melt temperature for each protein occurs in step 8 after which the melt shift is calculated in step 9. The final step 10 involves the summary of the proteins with significant stabilization or de-stabilization.

### Data exclusion, normalization and quantification

Raw abundance values reported by a proteomics search algorithm consist of the relative number of ions detected from a peptide homologous with an associated protein. Prior to beginning analysis by the TPP-TR pipeline, proteins can be excluded or filtered based on pre-determined quality control criteria. The purpose of the filtering is to address the technical variability that is present from the sample harvesting to LC-MS/MS analysis. One criterion used is whether a protein of interest is present in both data sets used to calculate the melt shift. In the event that a protein is observed in only one of the two conditions, the protein will be filtered from the analysis and will not be included in further analysis. This step is not unlike data preprocessing done for other types of quantitative proteomics studies to deal with the challenges of missing values from multiple MS-acquired datasets.

The raw untreated data from the proteomic analyses for each of the 12 data sets were analyzed to determine the total number of proteins that were present in each condition so that the level of exclusion could be quantified. We calculated the number of proteins that were present in each condition (i.e. mutant, WT), along with the number of proteins that were not present in the compared group. In the case of the Peck-Justice data sets, this analysis involved comparing the WT and *rpn5* and *pup2* mutant data sets from two biological replicates. The analysis done on the Perrin data sets compared the melt shift between the proteins in the PBMC and the whole blood data sets. The rodent organ data sets from Perrin were used by comparing the melt shift between the kidney and the liver data. While between tissue melt shifts were not specifically reported by Perrin et.al, other studies have recently reported similar types of analyses ^13^.

Figure 2 presents the large number of proteins for which melt temperature is only able to be calculated in one of the two data sets where the bar chart shows the large number of proteins that are present in only one of the two data sets being used to calculate a melt shift. This figure shows specifically that 7-32% of the total number of proteins would not be used in further analysis if an exclusion step were to be used in the pipeline. Upon further examination, it was confirmed (data not shown) that the reason for proteins being exclusive to only one of the two data sets (i.e. mutant or WT) was due to a generally low abundance for the protein of interest in the systems studied. These low abundance values have the impact of lowering the statistical values used to describe the total proteome or meltome (downstream analysis). The impact to the quantified meltome would be even more noticeable with the use of the mean function that is more sensitive to data skew. It is important to note as well that the use of newly reported mass spectrometry techniques like the use of a BOOST channel^14^ would impact this step in the analysis pipeline and whether proteins have sufficient abundance to be included in further analysis. Additionally, this clearly shows that efforts to increase the overall protein depth of coverage across samples is an important metric for this TPP / CETSA analysis as it is for global proteomics studies. Overall, the impact and reason for exclusion needs to be understood fully before considering use of this step in the data analysis pipeline.

**Figure 2.**
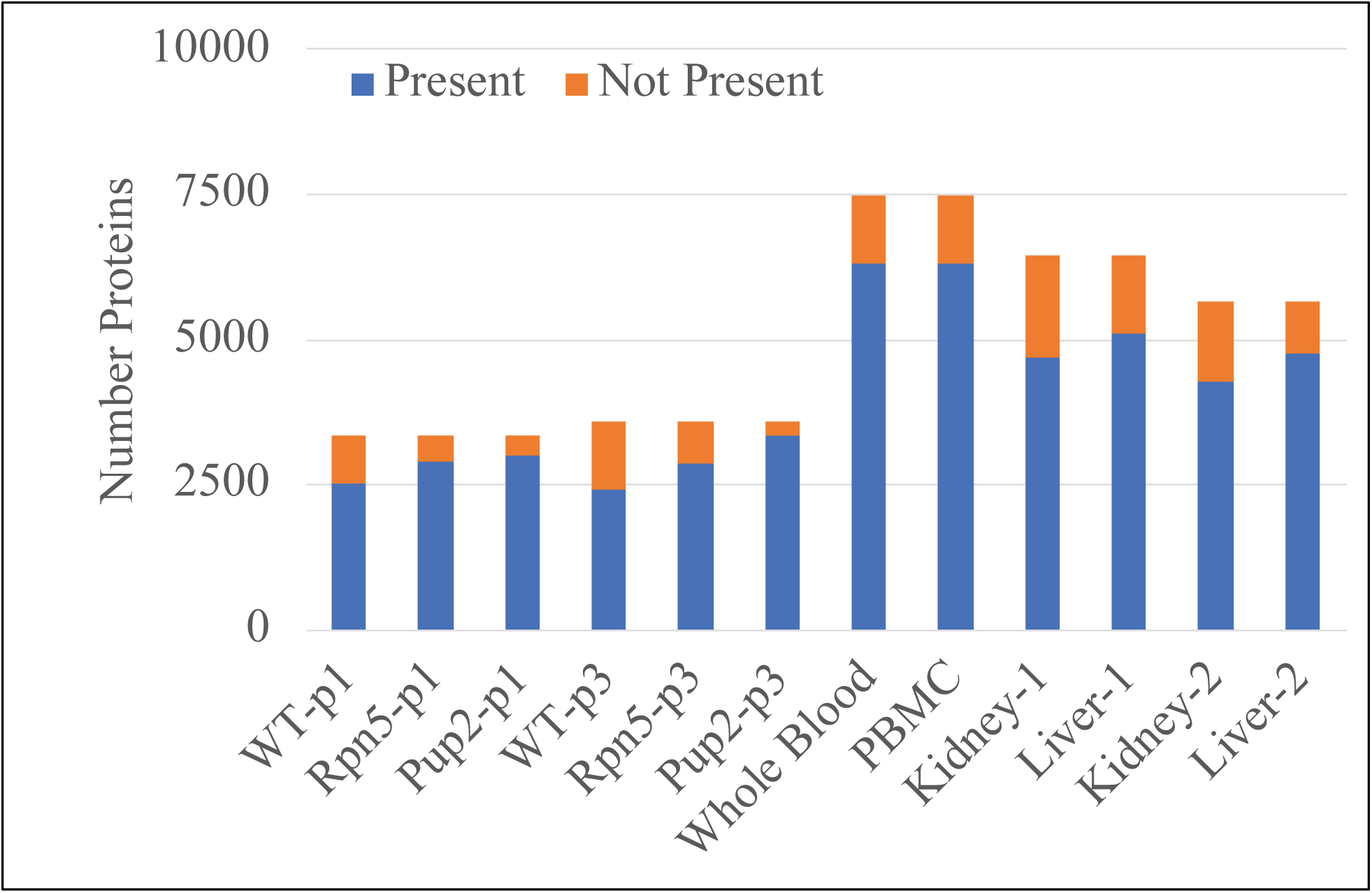
Determination of the number of quantitated proteins in each of the 12 experiments analyzed in this report. Each study or set of experiments that were compared in the data analysis are separated by the dashed lines. The total number of proteins that were observed in each group (i.e. p1 replicate from the Peck-Justice data set) is represented by the total height of the blue and orange bars. The number of proteins that were present in the corresponding data set are represented by the blue portion of the bar while the number of proteins that were not present in the corresponding data set are represented by the orange bar.

After low abundance proteins are excluded, all proteins abundance values at each temperature in the heat treatment are divided by the abundance observed at the lowest temperature in the heat treatment. This normalization step not only sets a reference of abundance to the lowest heat treatment temperature (or the theoretical max protein abundance) but it also converts protein abundance values to an equivalent scale so that they can be compared between different conditions. Results from abundance normalization for the Peck-Justice and Perrin data sets are shown in Figure 3 and Figure 4 respectively. The data in these sets of dot plots show not only the spread of abundance values that has been observed to occur at each temperature but the median bars in the plots demonstrate examples of the general departure from ideal sigmoidal shape that can occur. The sources of variability in a multiplexed workflow like this has previously been described to be due to a host of challenges ranging from technical differences to TMT label variation.^15-17^ Post normalization, abundance values across all proteins in each condition are then quantified statistically by use of mean or median functions. The calculated statistic and the corresponding proteome melt curves numerically describe the total abundance of proteins for a particular treatment or mutation. Curve fitting methodologies (described in the next section) are afterwards utilized to describe and predict the total protein abundance as a function of heat treatment temperature. Differences between actual and predicted protein abundance are used to calculate correction constants for each heat treatment temperature. The correction process adjusts the abundance of each protein at each temperature for any departures of the global meltome from expected melt behavior. Curve fitting, used a second time (and per the next section), is then used to describe the normalized abundance of each individual protein as a function of heat treatment temperature.

**Figure 3.**
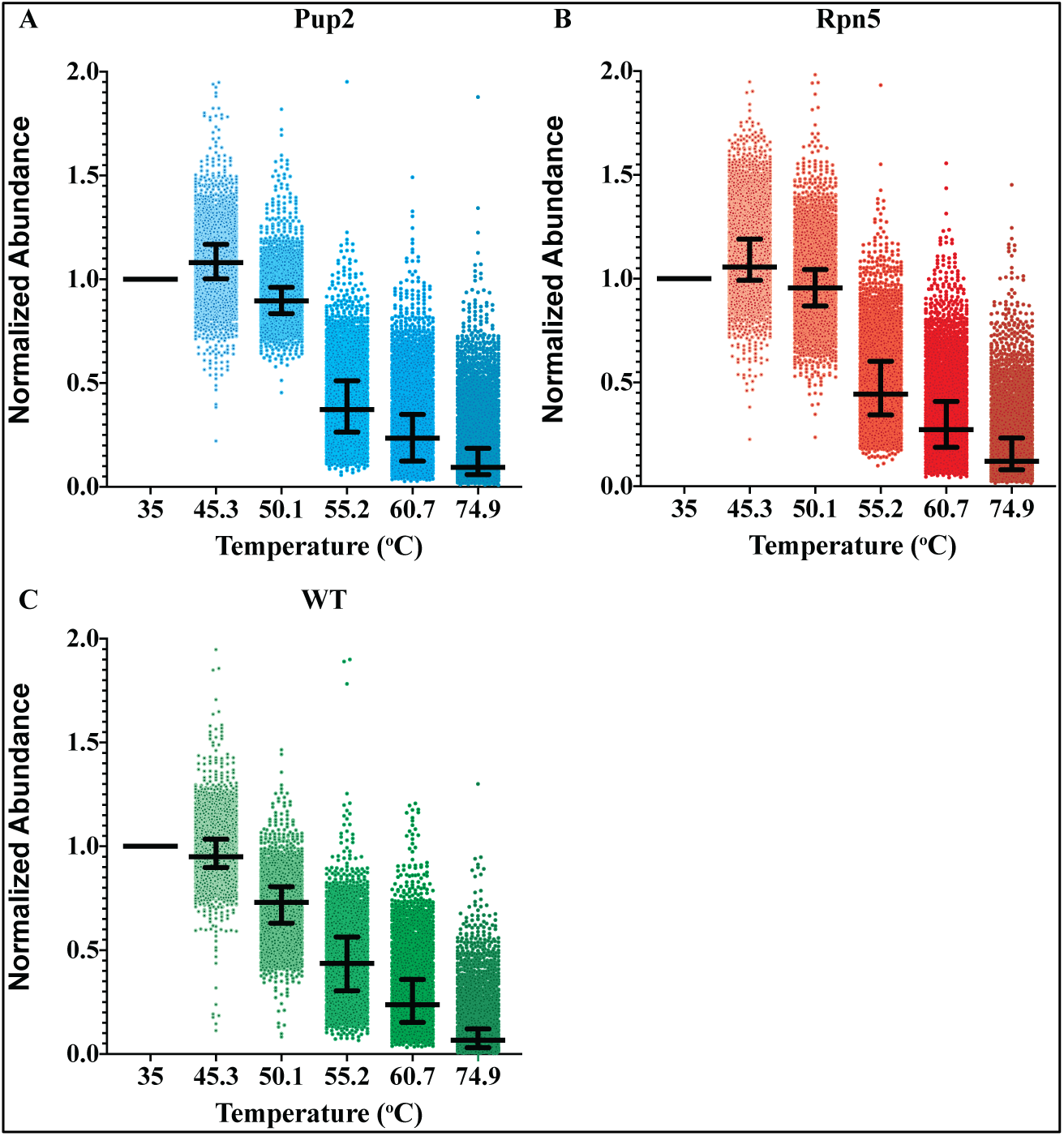
Dot pots of normalized abundance at each temperature relative to the abundance at the lowest temperature. The values are from the p1 and p3 data sets for Pup2 (A) Rpn5 (B) and WT (C) from the Peck-Justice data sets. Median and interquartile ranges are shown in the box and whiskers with the maximum axis value at 2 for easier viewing.

**Figure 4.**
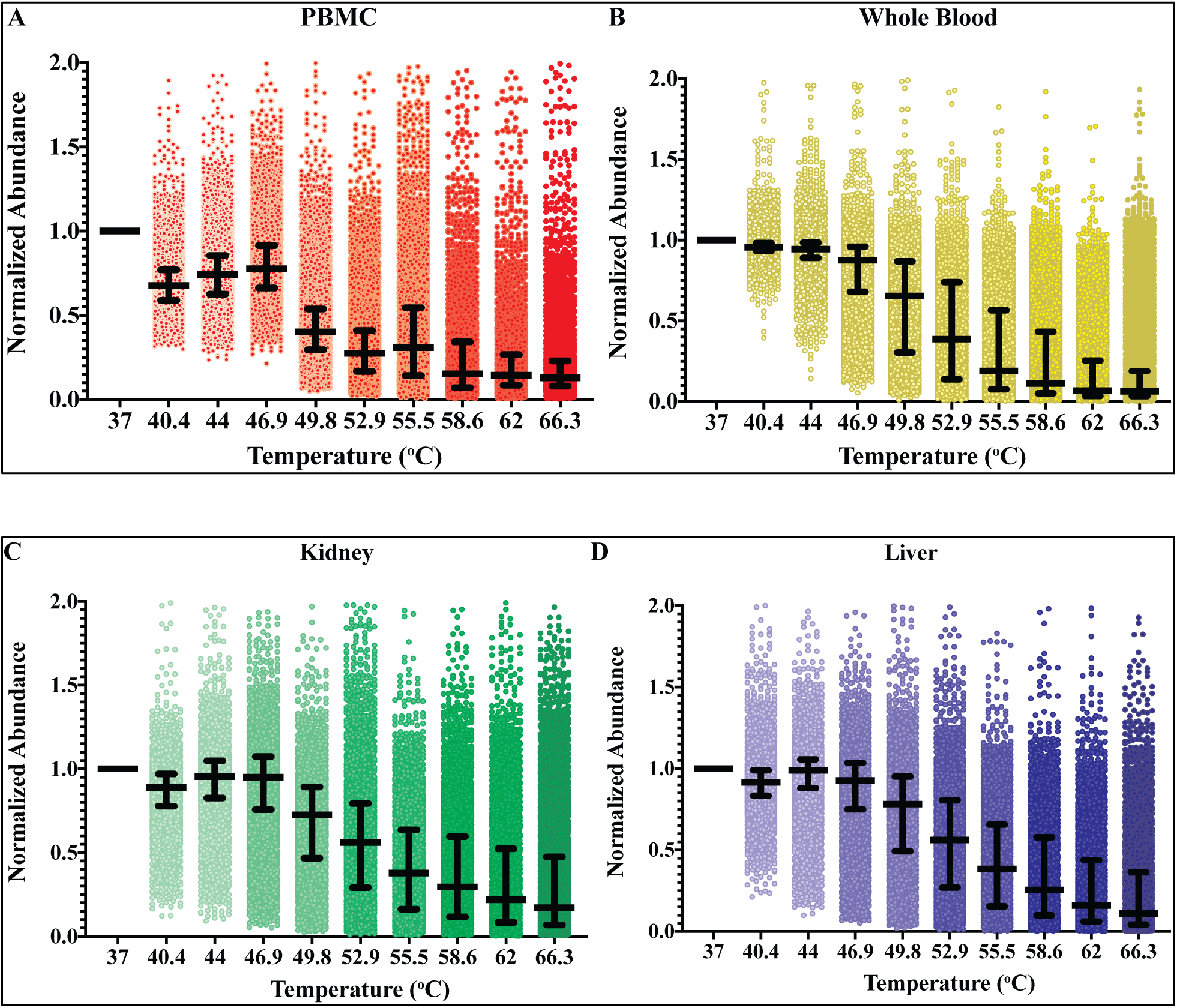
Dot plots of normalized abundance at each temperature relative to the abundance at the lowest temperature. The values are from the R2 data sets for PBMC (A) and human Whole Blood (B) Perrin data sets. The values are from the R1 and R2 data sets for (C) rat kidney, (D) rat liver Perrin data sets. Median and interquartile ranges are shown in the box and whiskers with the maximum axis value at 2 for easier viewing.

### Curve fitting

A melt curve with its sigmoidal shape can be described mathematically by a logistic expression. Two non-linear equations were used in our evaluation to determine which is optimal for TPP / CETSA studies; a three-parameter log fit (3PL) and a four-parameter log fit (4PL). The 3PL equation uses three calculated constants a, b and Pl to describe the abundance as a function of temperature, T whereas the 4PL uses an extra constant to describe the variability. The 4PL constants a, b, c and d are equal to the slope at the inflection point, the inflection point, lower plateau and maximum plateau respectively. The normalized abundance vs. temperature for two proteins in the Peck-Justice data set are shown in Figure 5 and these two curves provide insight into the impact of fit equation on the melt curve. First, the curves fit for the two selected proteins are just below and above our commonly used cutoff criteria of R^2^ (0.95) depending on which equation was used. In the case where the 3PL is used, the goodness of fit is below this example criteria whereas the 4PL fit results in a better fit that would be quantified as significant by the workflow. Another point from this analysis is that the fitting equation can also contribute to the curvature of the melt plot. In the case of (B), the curvature of melt is much steeper for the 4PL than for the 3PL fit. The steeper 4PL curve has a more clearly defined inflection (point on the line where curvature changes direction) than the 3PL fit and would have a more defined melt temperature if the inflection point definition were to be used. The 3PL fit, however, has a higher top plateau (crosses the y-axis at 1instead of 0.9) and has a shallower curve down the heat treatment. The impact of the more “stretched out” melt curve could affect the defined melt temperature depending on where the lower plateau of the 3PL curve levels off (greater than 75°C in panel B).

**Figure 5.**
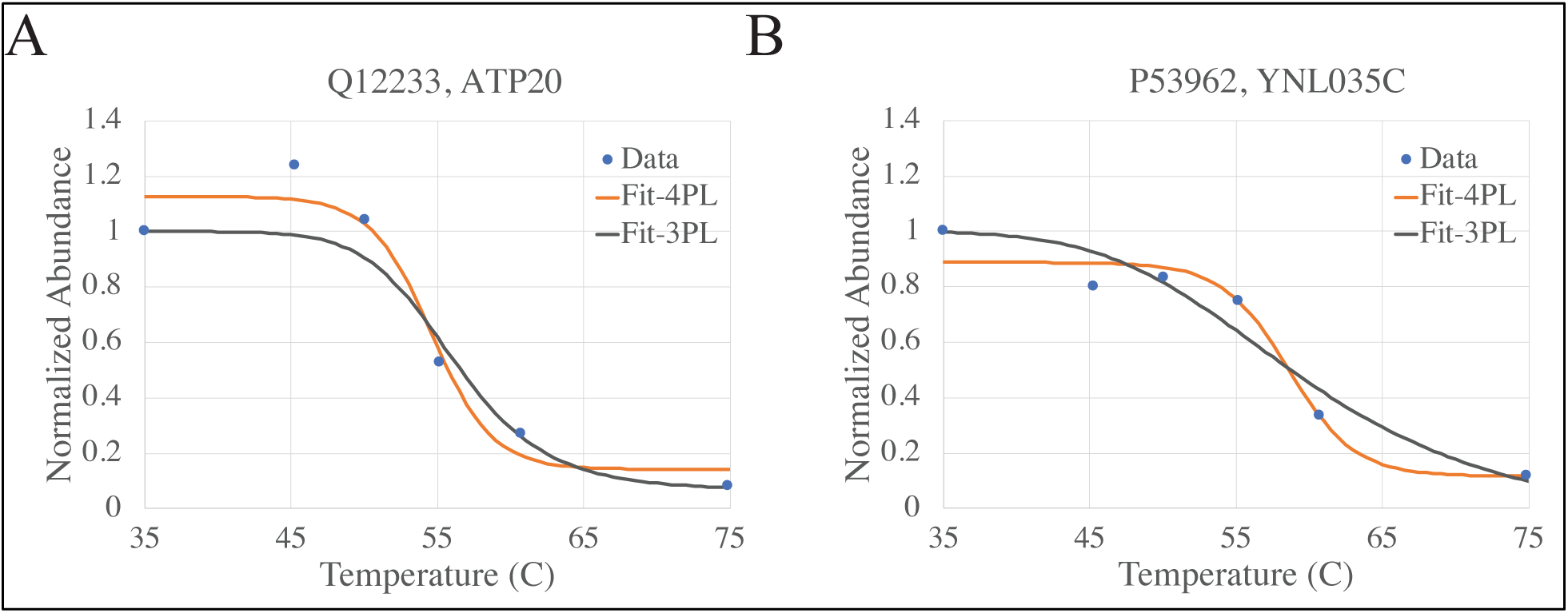
Comparison of 3PL Fit and 4PL Fit using normalized abundance at each temperature relative to the abundance at the lowest temperature. Data presented is from the Pup2-p1 data set. The 3PL and 4PL equations were used to fit the normalized abundance vs. temperature for each protein reported. (A) Q12233 which has an R2 of 0.92 for the 3PL and 0.96 for the 4PL fit. (B) P53962 which has an R2 of 0.93 for the 3PL and 0.96 for the 4PL fit.

The overall ability for a mathematical expression to describe the observed variability in a series of data points is commonly done using the coefficient of determination, R^2^. This coefficient used in linear systems equals the percentage of variability that is described by the independent variable. The sum of the residual squared error and regression squared error in a nonlinear or logistic system, on the other hand, does not necessarily equal the sum of squared total error and therefore the R^2^ can lie outside of the range of 0 to 1. The limitation makes the determination coefficient a poor measurement of fit for a nonlinear model. ^18-20^ One measurement of fit that was proposed ^21^ and evaluated ^19^ as a more suitable comparator for nonlinear fit than R^2^ is the Bayesian Information Criterion (BIC). The BIC is a quantitative evaluation of fit where more negative values indicate a more optimal regression between conditions. The WT-p1 data set from the Peck-Justice experiments was fit using both of the previously described 3PL and 4PL equations and the quality of fit was quantified using both the BIC and R^2^. The results in Figure 6 show that the 4PL fit provides models with comparatively higher R^2^ than the 3PL fit. While results shown in this figure indicate that the BIC is slightly better for the 3PL, the median value was lower for the 4PL. More work needs to be done to understand the meaning of the BIC especially as it relates to comparing results between different curve fitting methods.

**Figure 6.**
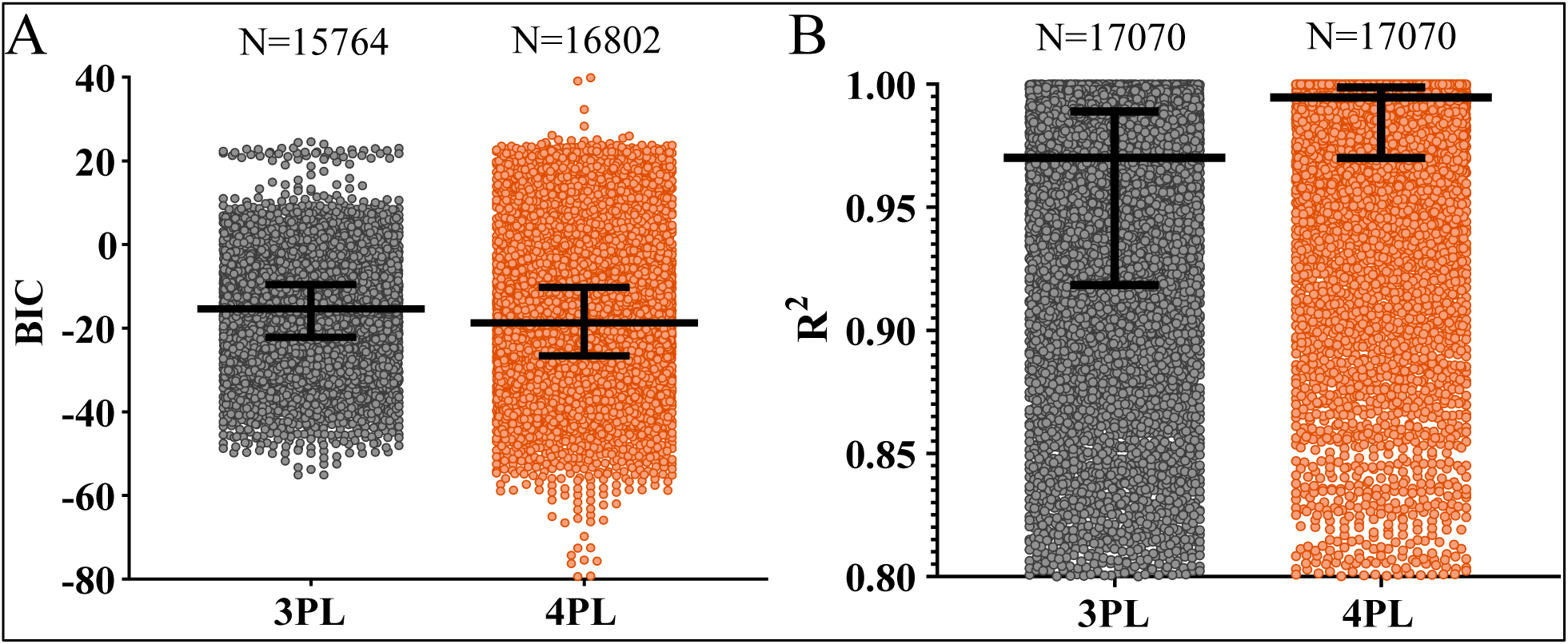
BIC and R2 for the two fit methods using all data from Peck-Justice data sets. Medians along with interquartile ranges are shown in the box and whisker plots. A portion of the points are shown in each data set; some points were outside of the y-axis range. Statistical analyses utilized unpaired t-test with Welch’s correction (A) The figure shows the BIC for the two fitting methods where medians are -15.4 and -18.7 for 3PL and 4PL, respectively. Maximum values are 24.6 and 228.4 for 3PL and 4PL respectively, while minimum values are -55.1 and -79.4 respectively. There is a statistically significant difference between the two data sets with p < 0.0001. The BIC could not be calculated for all conditions based on the curve shape thus the difference in N for the two methods. (B) The figure shows the R^2^ for the two fitting methods where medians are 0.970 and 0.995 for 3PL and 4PL respectively. Maximum values are 1.00 for both methods and minimum values are –3.383 and -3.614, respectively. There is a statistically significant difference between the two data outputs with p < 0.0001.

After the melt curves are described by their respective equations in the TPP-TR program, a single melt curve from one of the two conditions is used to calculate the correction factor for all conditions (Fig. 1, Step 5). The curve and corresponding condition with the best fit, as measured by the R^2^, is the condition that is used to calculate the correction constant for both conditions. An example of a result from this TPP normalization is shown in Figure 7 for the Rpn5 protein (Uniprot accession: Q12250) using the Rpn5-p1 data set. These plots show how the melt curve changes as a result of the correction step. The purple (A) and yellow (B) data points for each of the two plots show the normalized abundance for each protein at each temperature in the WT and mutant data sets respectively. The green (A) and red (B) data points are the normalized abundance values for the Rpn5 protein prior to correction while the black (A) and orange (B) points are the values post correction. The 3 parameter fits to the corrected points in A and B are shown in green and blue respectively and show no significant correction in the case of the WT data set. The curve shift for the 3-parameter fit that resulted from the correction to the mutant data set, on the other hand, was more significant than the WT data set. These data are informative in a couple ways. First, it is important to note that correction can help to abrogate overall differences in curve shape that likely result from technical variation between samples making it key for direct sample comparison Second, the datasets included in the analysis pipeline simultaneously is a key consideration and should be limited to things that are directly being compared.

**Figure 7.**
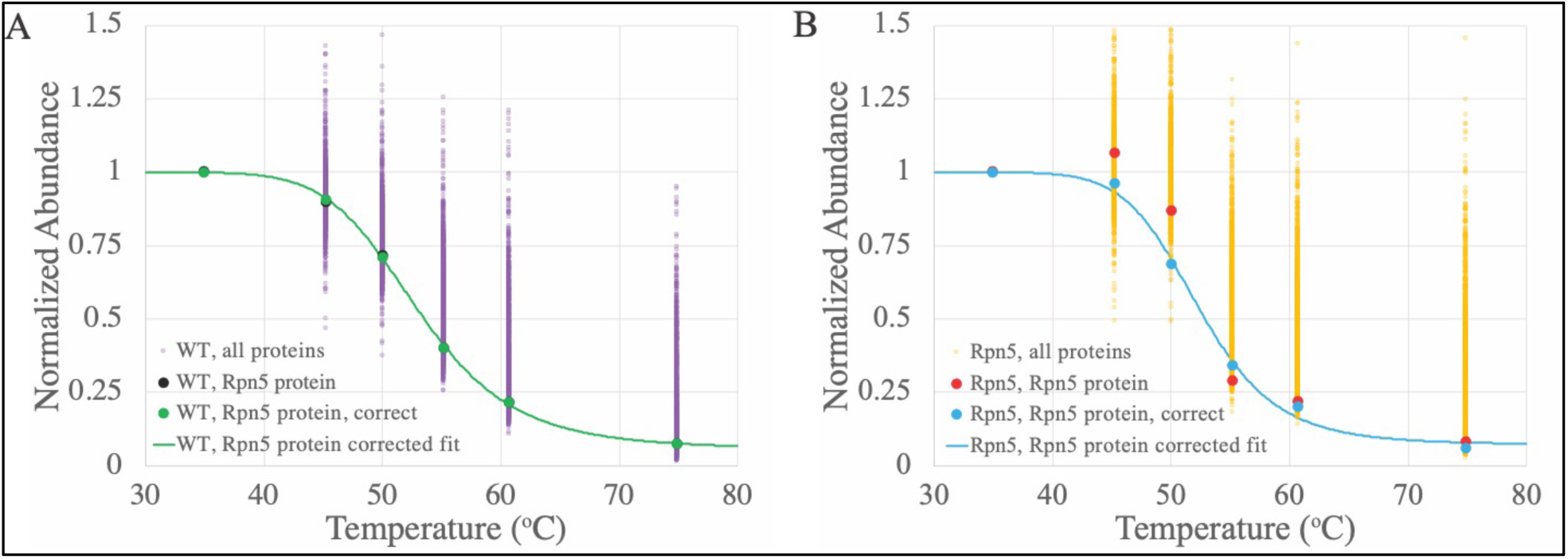
Global normalized abundance at each temperature relative to the abundance at the lowest temperature for both p1 data sets for (A) WT and (B) mutant Rpn5. Individual purple or yellow data points are for the individual proteins in each data set. In the case of panel (A), the black and green dots are the Rpn5 protein normalized abundance values in the WT-p1 data set, before and after data correction respectively. In the case of panel (B), the red and blue dots are the Rpn5 protein normalized abundance values in the Rpn5-p1 data set, before and after data correction respectively. The green line in panel (A) is the best fit line for Rpn5 protein in the WT-p1 data set while the blue line in panel (B) shows the best fit line for the Rpn5 protein the Rpn5-p1 data set.

### Melt temperature calculation and exclusion

The melting point, T_m_, of any protein is defined as the temperature at which a protein unfolds from its native state. Due to the fact that all proteins in solution do not unfold *en bloc*, the melt point is often defined as a transition point in one of two ways. While some sources have defined the melt as the point at which 50% of the protein remains folded ^22^, others consider this transition to be the inflection point in a melt curve ^23, 24^. In order to study the impact of melt definition on the analysis output, the normalized abundance values for the proteins at the highest treatment temperature across all experiments were plotted (Figure 8). Each of the values in these plots correspond to the bottom plateau of each melt curve and should ideally cross at the value of 0 if all protein is denatured and is separated from the liquid at the highest temperature. These plots show that despite the median abundance being near zero, there are a significant number of proteins in each data set with normalized abundance above 0.5.

**Figure 8.**
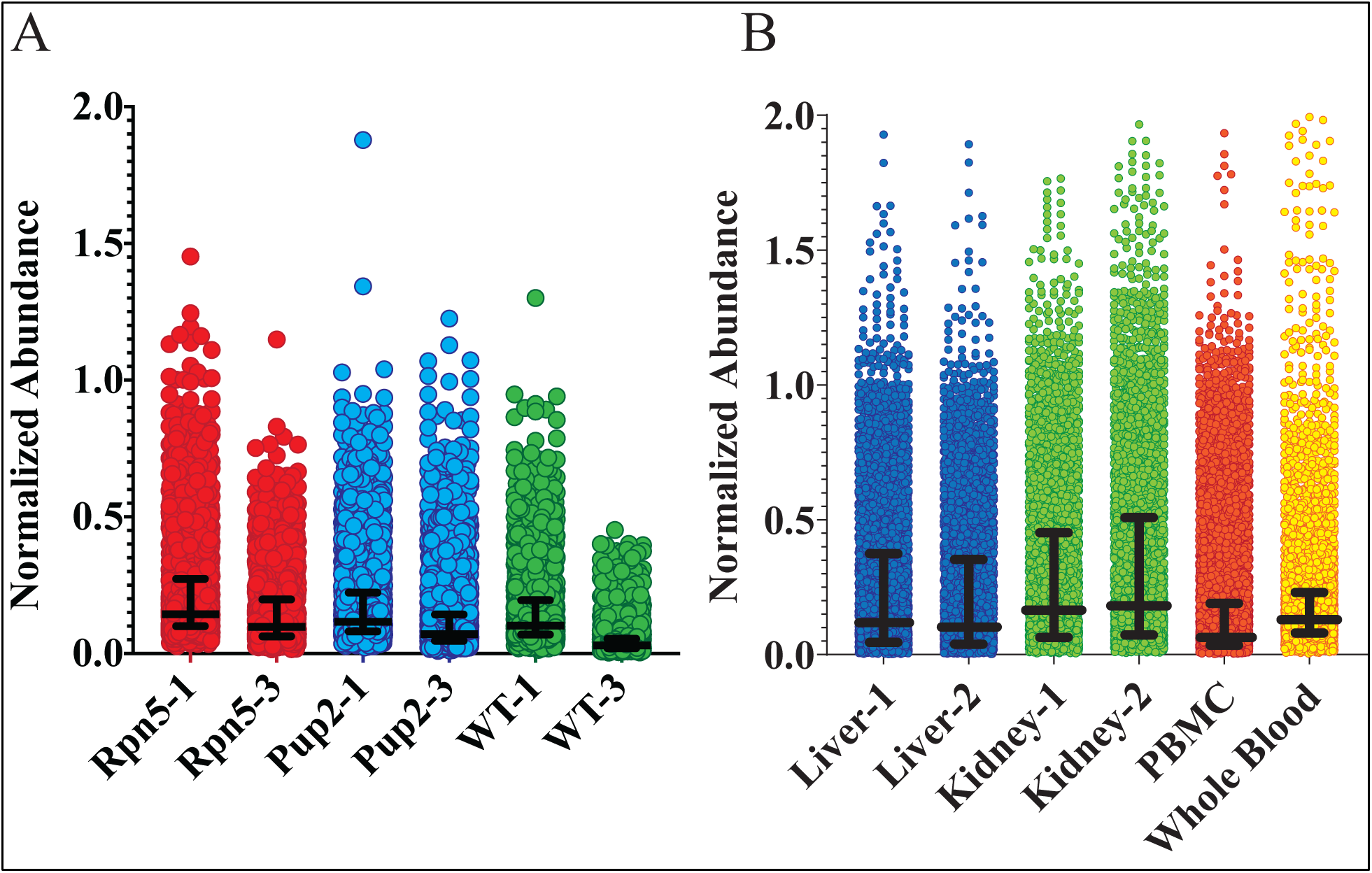
(A) Comparison of normalized abundance at each temperature relative to the abundance at the lowest temperature for each of the highest temperatures in each study, Peck-Justice data sets. Note that the y-axis range has been decreased in order to make the box plots viewable. (B) Comparison of normalized abundance at each temperature relative to the abundance at the lowest temperature for each of the highest temperatures in each study, Perrin data sets. Note that the y-axis range has been decreased in order to make the box plots viewable

Two of the melt curves from the TPP-TR pipeline analysis are also plotted in Figure 9 and the example data in (A) reinforces why some of the proteins do not have calculated melt temperatures. The melt curve for protein PRDX4 (Uniprot accession: Q13162) in the “mutant” condition has a lower plateau higher than 0.5 and consequently does not have a defined melt temperature with the 50% definition. A lack of melt temperature from one of the two conditions (“WT” or “mutant”) results in no melt shift calculation, which reduces the overall number of comparative values being reported. In the case of Figure 9 (B), the definition of the melt being equal to 0.5 also has a significant impact on the calculated shift. The “mutant” curve for RNF170 (Uniprot accession: Q96K19) crosses the 50% denatured point at 53°C while the inflection point of this same curve is closer to 51%. The melt shift based on a definition of where the curves equal 0.5 can result in a different shift than if the definition is based on the inflection point of the curves.

**Figure 9.**
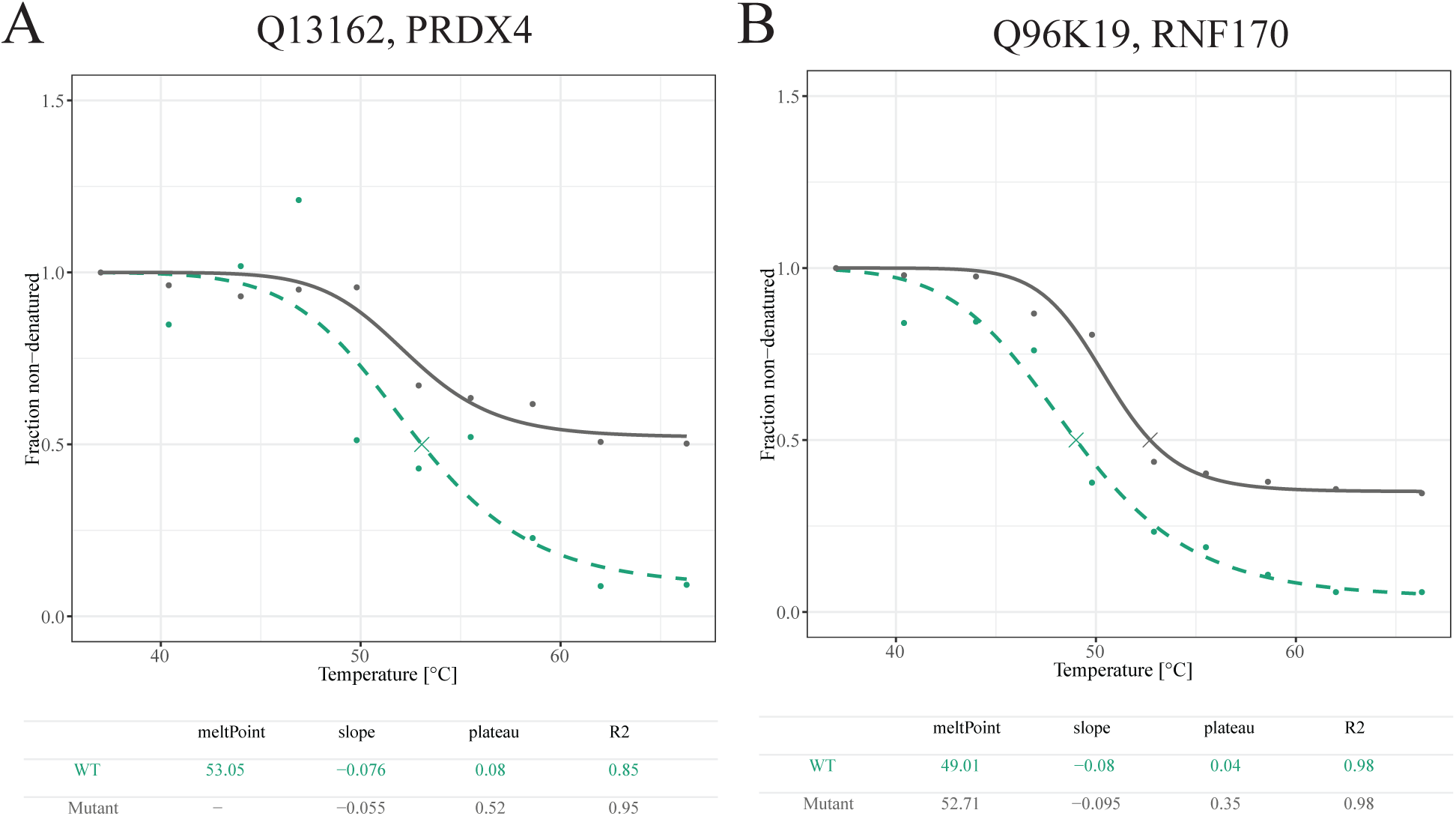
Examples of melt curves from TPP-TR method using Perrin data set (A and B) where calculated melt temperature is limited by definition of melt (i.e. temperature at which 50% soluble protein is lost). (A) is PRDX4 or Q13162 and (B) is RNF170 or Q96K19. The curves generated using this method either did not generate melt temperatures for all of the conditions due to curves not crossing 0.5 (A) or had a calculated melt shift significantly impacted by the definition of the melt temperature (B).

### Melt shift calculation and stability summary

Each of the data sets collected by Peck-Justice and Perrin were analyzed using both the Childs *et al*. TPP-TR workflow along with our proposed workflow in order to understand the relative quantity of significantly stabilized and destabilized proteins. Our assessment used the same goodness of fit criteria (melt curve R^2^ of 0.95) and melt shift significance cutoff (2 standard deviations from mean, a 95% confidence interval) for both pipelines. In the case of the Pup2 and Rpn5 data sets, each curve was compared with its corresponding WT data set in order to calculate melt shifts. In the case of the Perrin human blood data set, PBMC values were used as the experimental condition while whole blood results were used as the control data set. This analysis was not carried out in the original published report but was used in our analysis for comparison to illustrate potential shifts in melt between a specific fraction of blood and the bulk of human blood matrix. Additionally, the rodent organ data analysis was done by comparing each of the kidney with the liver data sets with the goal of showing comparative protein stabilization/destabilization between each organ and the liver.

Once each data set was evaluated using the two workflows, the number of proteins with significant melt shifts results were compared using R-based Venn diagrams. The first comparison (Figure 10) illustrates the amount of overlap between the two analysis outputs relative to the total number of proteins in each data set. These diagrams show that while 40-78 proteins were shared between the two methods, there were an even greater number of proteins in each case that were not observed as significant by the other corresponding analysis pipeline. This result reflects the strong sensitivity of the analysis output to the type of steps that are used in the analysis.

**Figure 10.**
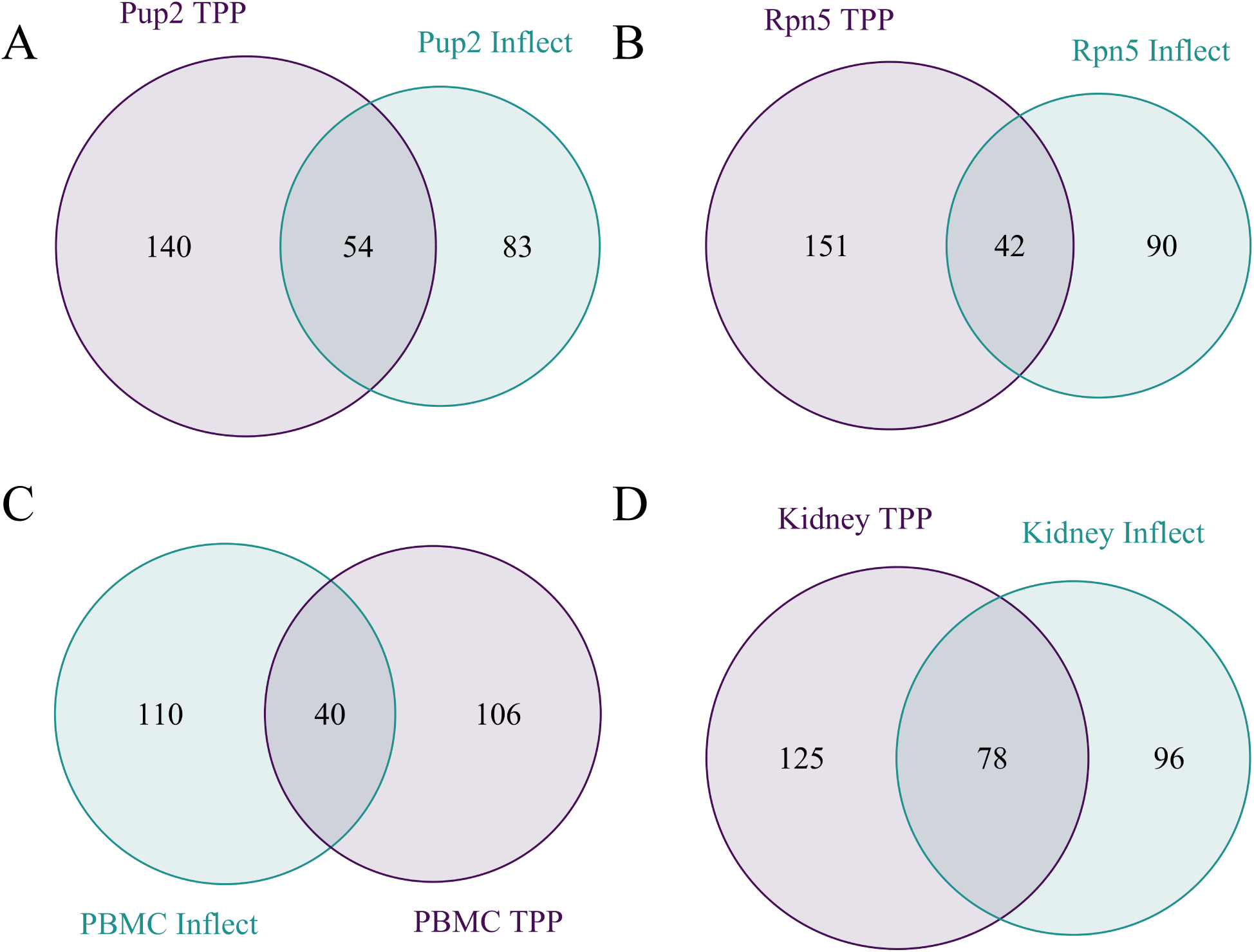
Melt shift evaluations using each workflow, TPP-TR and our workflow. Comparison of significant proteins observed between the two pipelines. The Peck-Justice data sets for Pup2 and Rpn5 relative to WT are shown in (A) and (B). The Perrin data sets are shown in (C) through (D). Human PBMC relative to human whole blood are shown in (C) while rat kidney relative to rat liver are shown in (D).

While there were a large number of proteins that were found to be uniquely significant in our workflow, we also found many of these proteins were relevant to the study or question being asked in the original data set. In the case of the Pup2 data set, Pre1, a component of the 26S proteasome, was reported as destabilized in our data set but was not found to be significant using the same criteria for significance in the TPP-TR workflow. The TPP-TR workflow neglected to find this shift as significant due to the fact that the melt shift for the wild type condition was just below the fit quality criteria of 0.95. This protein is of interest due to the fact that the strain used in the reported experiment leveraged a mutation to the Pup2 gene, another component of the 26S proteasome and thereby a potential protein-protein interaction partner.^5^ The negative shift in the melt temperature indicates that the *pup2* mutation resulted in a destabilization of the Pre1 protein with other proteins, potentially those in the 26S proteasome. Other proteasome or ubiquitin related proteins that were observed as significant in our workflow are shown in Table 1 and the associated melt curves for some of these proteins are displayed in Figure 11. A similar trend was observed for the Rpn5 data sets where a significant number of proteasome subunits were observed with significant melt shifts in our workflow only, which was not uncovered in the initial published study. The proteins of interest are shown in Table 2 while some of the melt shifts from these Rpn5 proteins are shown in Figure 12. We observed that in the case of more than one of the “WT” data sets, the melt curves had a higher than average inflection point which suggests high thermal stability in WT cells. It has already been shown by others that the proteasome and ubiquitin have higher melt temperatures than the average protein and therefore would have implied greater thermal stability ^25, 26^. It is possible that proteins with high intrinsic thermal stability may represent dataset outliers; however, to facilitate the development of TPP analysis methods that facilitate biological discovery it is important that optimized analysis pipelines consider proteins with a wide array of biophysical properties.

**Table 1.**
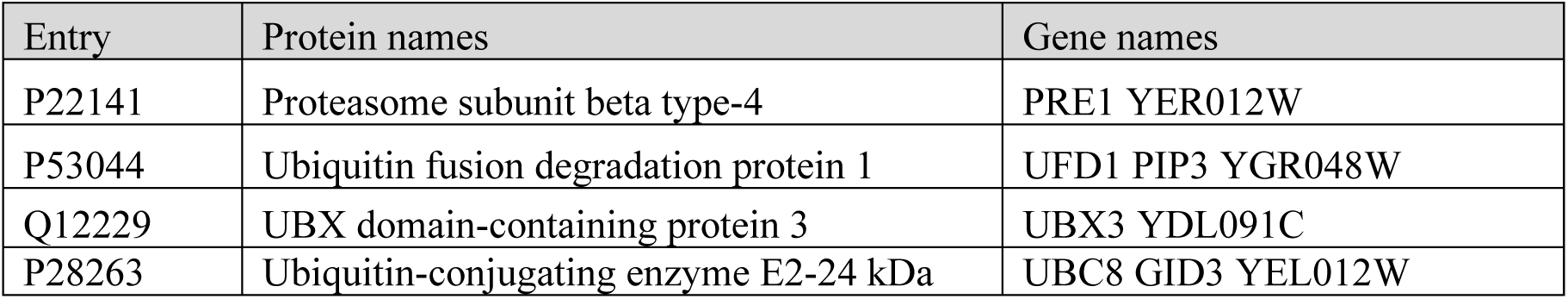
Summary of proteasome related proteins (based on information from Uniprot) that had significant temperature shifts from the Pup2 data sets using our workflow but were not observed to be significant in the TPP-TR workflow.

| Entry | Protein names | Gene names |
| --- | --- | --- |
| P22141 | Proteasome subunit beta type-4 | PRE1 YER012W |
| P53044 | Ubiquitin fusion degradation protein 1 | UFD1 PIP3 YGR048W |
| Q12229 | UBX domain-containing protein 3 | UBX3 YDL091C |
| P28263 | Ubiquitin-conjugating enzyme E2-24 kDa | UBC8 GID3 YEL012W |

**Table 2.**
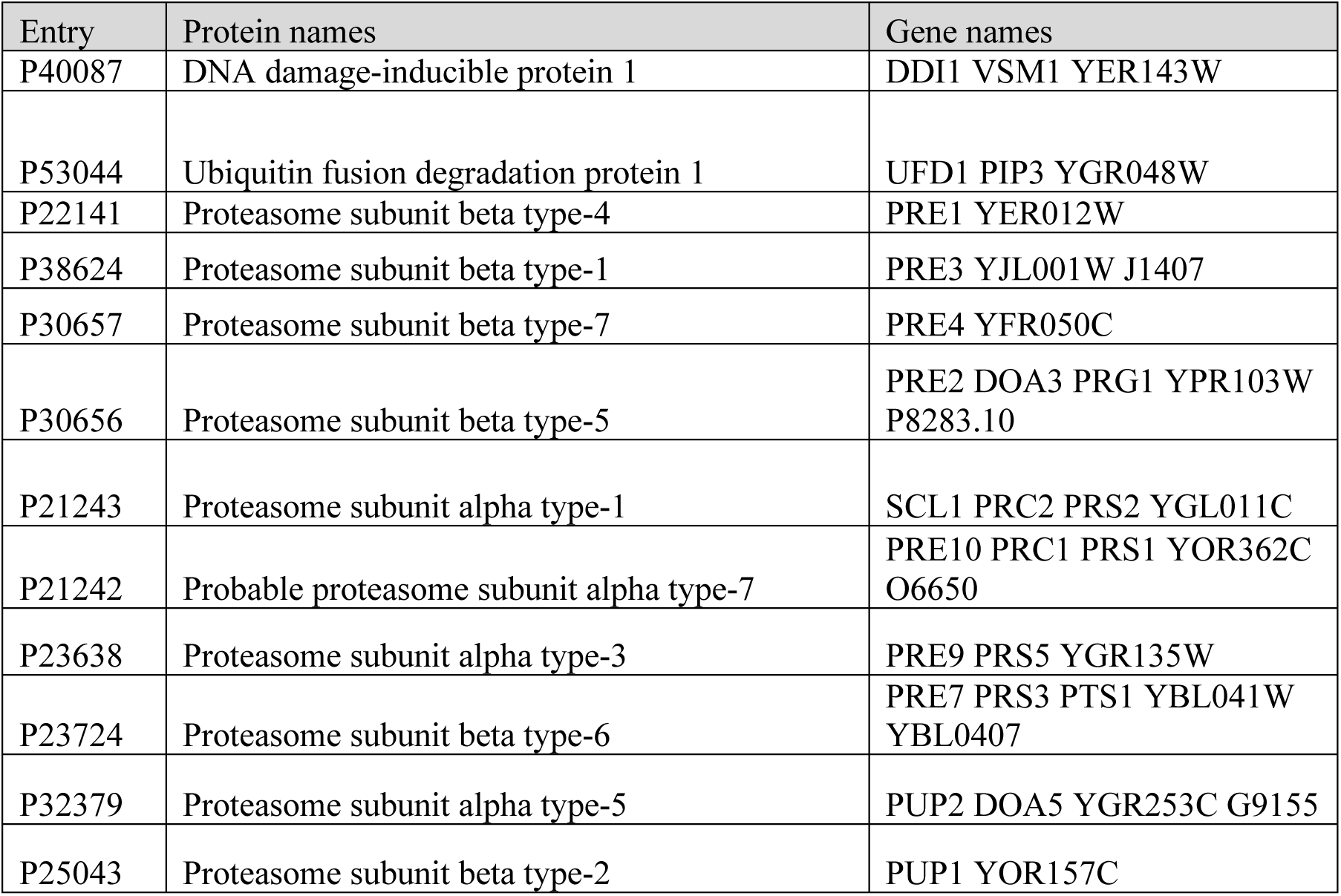
Summary of proteasome related proteins (based on information from Uniprot) that had significant temperature shifts from the Rpn5 data sets using our workflow but were not observed to be significant in the TPP-TR workflow.

**Figure 11.**
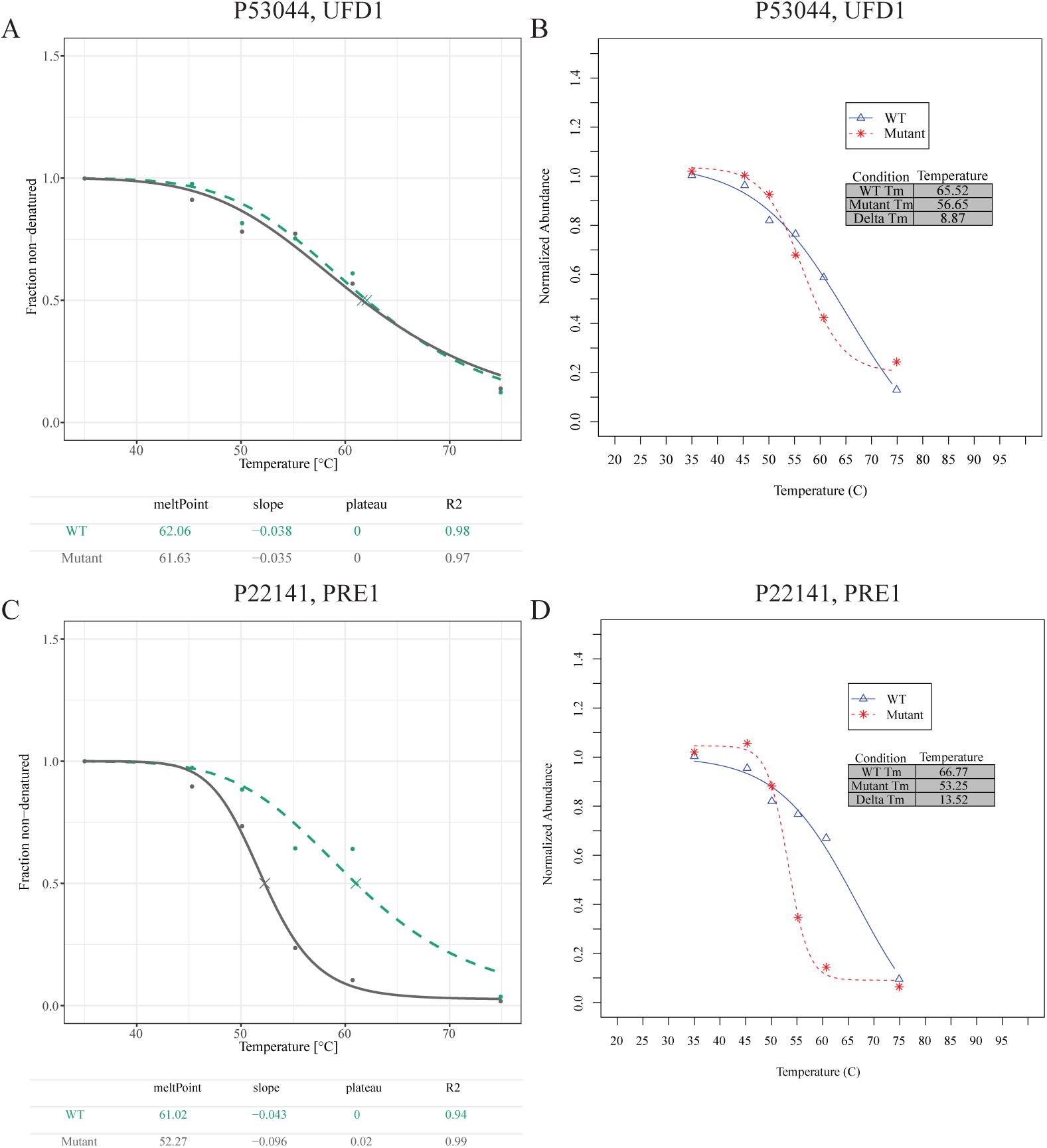
Pup2 data set melt shifts from Peck-Justice data sets that were reported significant in our workflow but not significant in the TPP-TR workflow. Melt curves generated from the Ubiquitin fusion degradation protein, Ufd1 using (A) TPP-TR and (B) our workflow. Melt curves generated from the Proteasome subunit beta type-4, PRE1 using (C) TPP-TR and (D) our workflow.

**Figure 12.**
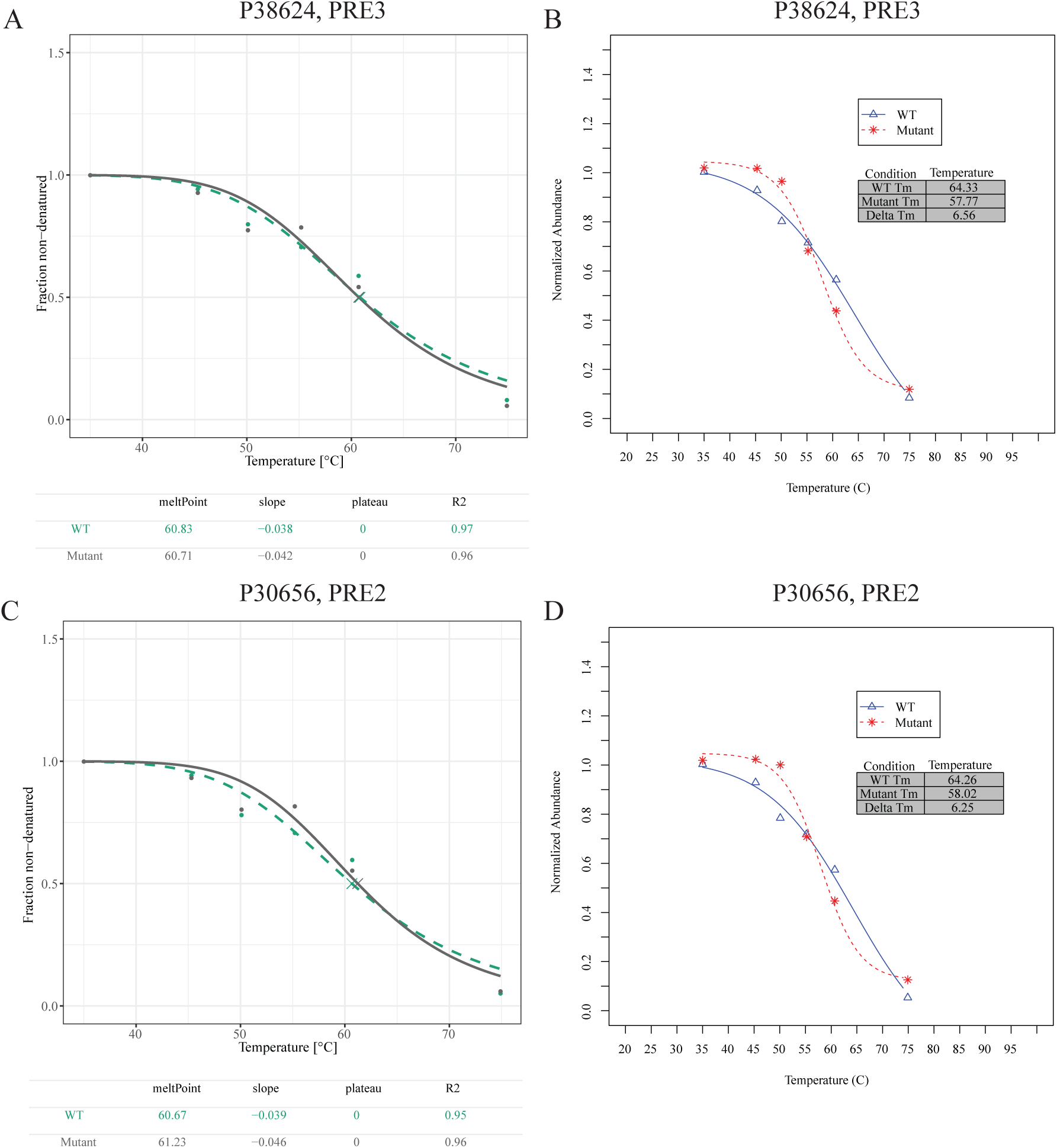
Rpn5 data set melt shifts from Peck-Justice data sets that were reported significant in our workflow but not significant in the TPP-TR workflow. Melt curves generated from the Proteasome subunit beta type-1, PRE3 using (A) TPP-TR and (B) our workflow. Melt curves generated from the Proteasome subunit beta type-5, PRE2 using (C) TPP-TR and (D) our workflow.

PBMCs are by definition the fraction of blood that play a significant role in the immune response and are enriched in T-cells, B-cells, NK cells and monocytes ^27^. Proteins from the PBMC vs. whole blood data that were observed as significant in our workflow but not significant in the TPP-TR workflow while also being relevant to a hematological or immunological function are highlighted in Table 3. The melt curves for three of these proteins with their corresponding comparisons between the two data analysis pipelines are shown in Figure 13. *Rattus norvegicus* TPP experiments that were executed by Perrin *et al*. were also examined for relevance to the organ being studied. In the case of the kidney data that was compared with the liver data set, there were several proteins that were unique to our workflow output. Many of these proteins (Table 4) have reported specificity for the kidney based on their functional annotations found within the Uniprot database ^28^ and thus suggest biological relevance for proteins found by our workflow. Examples of compared melt curves are shown in Figure 14 and each of these curves emphasize unique causes for results not being observed as significant in the TPP-TR workflow.

**Table 3.** Summary of proteins related to blood or leukocytes or expressed in blood cells (based on information from Uniprot) that had significant temperature shifts from the Perrin PBMC vs whole blood data sets using our workflow but were not observed to be significant in the TPP-TR workflow.

| Entry | Protein names | Gene names |
| --- | --- | --- |
| Q96EK5 | KIF-binding protein | KIFBP KBP KIAA1279 KIF1BP |
| O14672 | Disintegrin and metalloproteinase domain-containing protein 10 | ADAM10 KUZ MADM |
| P05556 | Integrin beta-1 | ITGB1 FNRB MDF2 MSK12 |
| P78325 | Disintegrin and metalloproteinase domain-containing protein 8 | ADAM8 MS2 |
| Q96LC7 | Sialic acid-binding Ig-like lectin 10 | SIGLEC10 SLG2<br>UNQ477/PRO940 |
| Q8WU39 | Marginal zone B- and B1-cell-specific protein | MZB1 MEDA7 PACAP HSPC190 |
| Q9NR28 | Diablo homolog, mitochondrial | DIABLO SMAC |
| Q9NP99 | Triggering receptor expressed on myeloid cells 1 | TREM1 |
| Q15631 | Translin | TSN |
| O76031 | ATP-dependent Clp protease ATP-binding subunit clpX-like, mitochondrial | CLPX |
| O95232 | Luc7-like protein 3 | LUC7L3 CREAP1 CROP O48 |
| Q68CP4 | Heparan-alpha-glucosaminide N-acetyltransferase | HGSNAT TMEM76 |

**Table 4.**
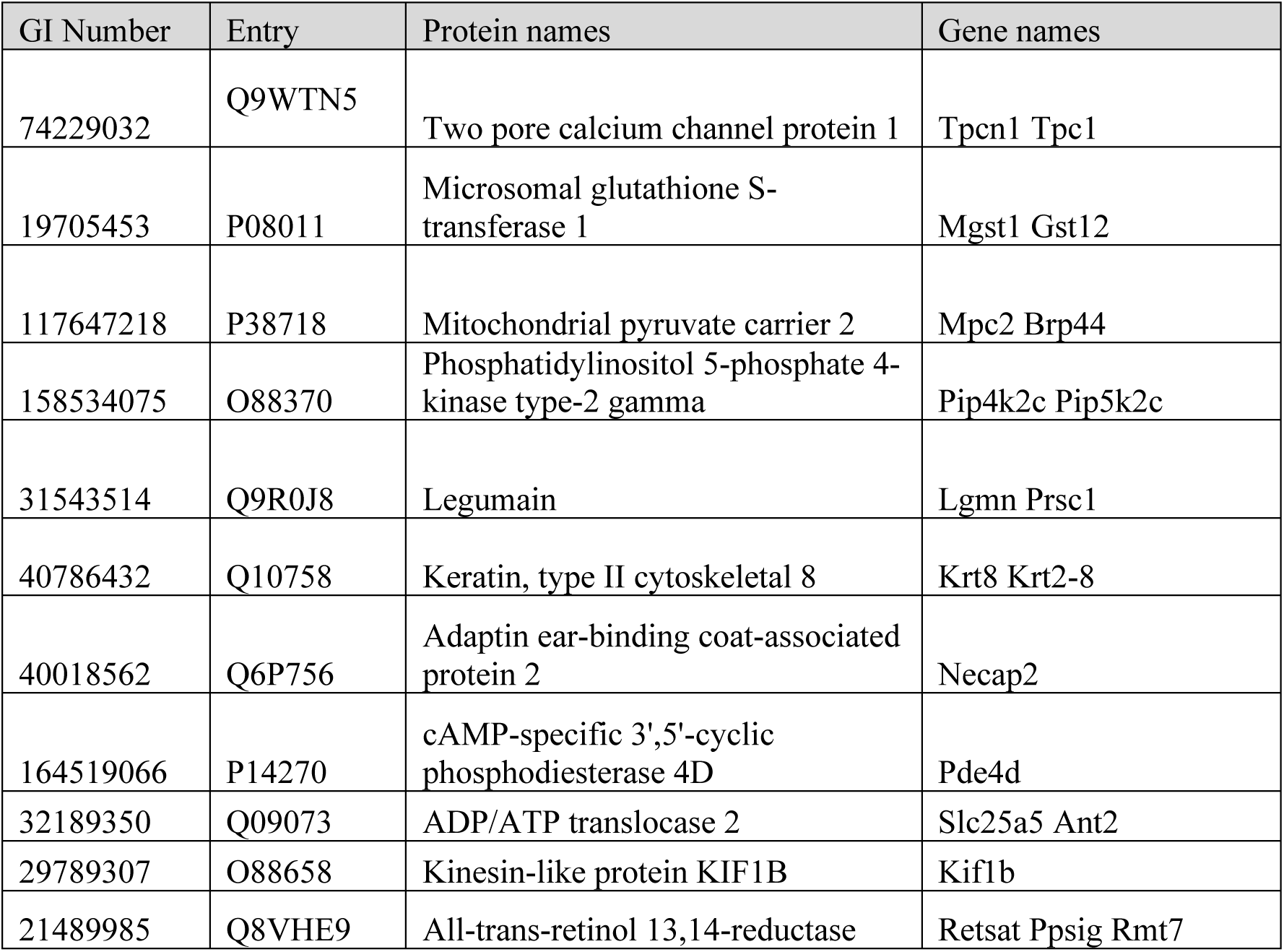
Summary of proteins related to kidney function (based on information from Uniprot) that had significant temperature shifts from the Perrin rat kidney vs liver data sets using our workflow but were not observed to be significant in the TPP-TR workflow.

**Figure 13.**
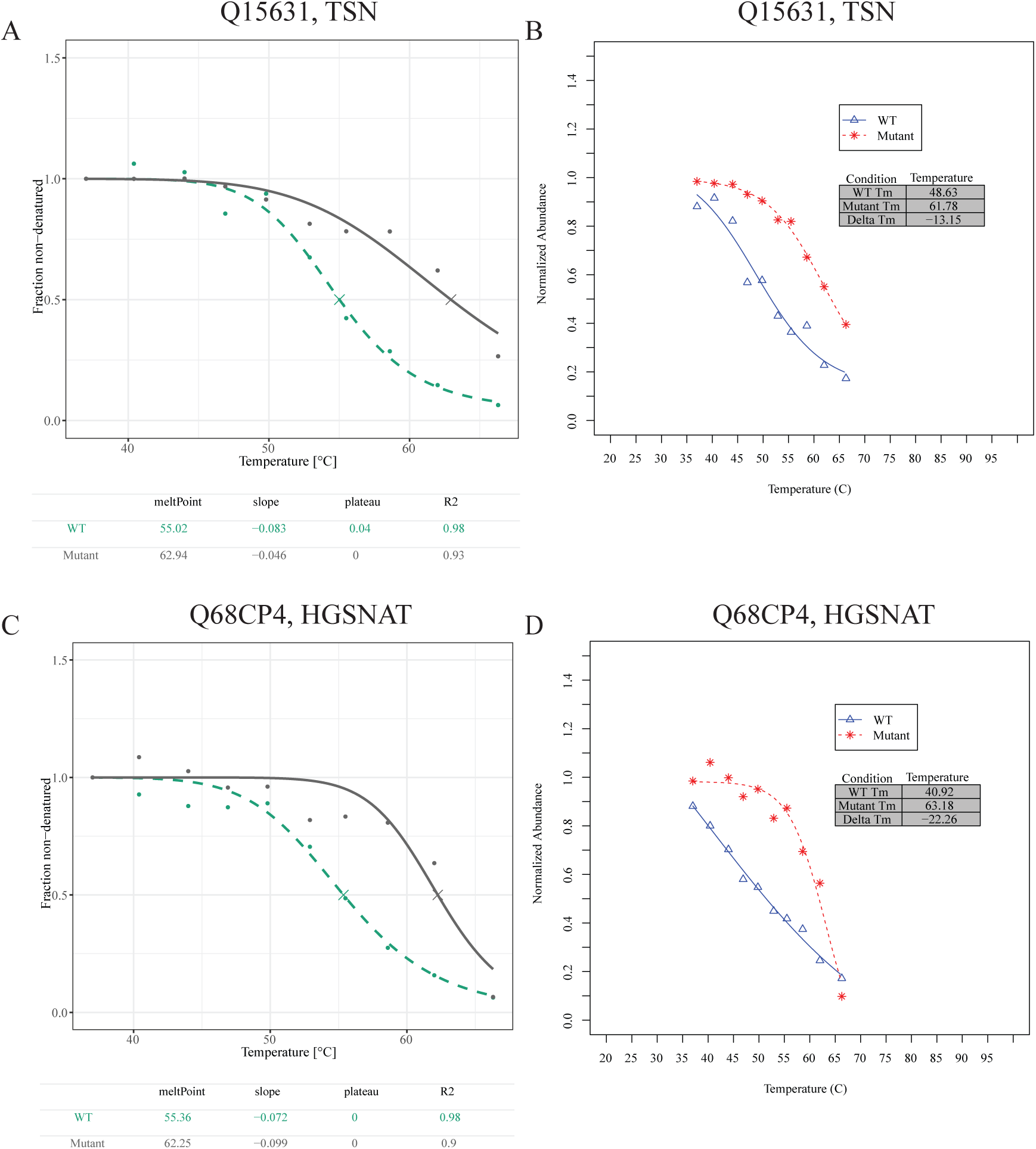
Example human PBMC vs. whole blood melt shifts from Perrin data sets that were reported significant in our workflow but not significant in the TPP-TR workflow. Melt curves generated from the Translin, TSN using (A) TPP-TR and (B) our workflow. Melt curves generated from the Heparan-alpha-glucosaminide N-acetyltransferase, HGSNAT using (C) TPP-TR and (D) our workflow

**Figure 14.**
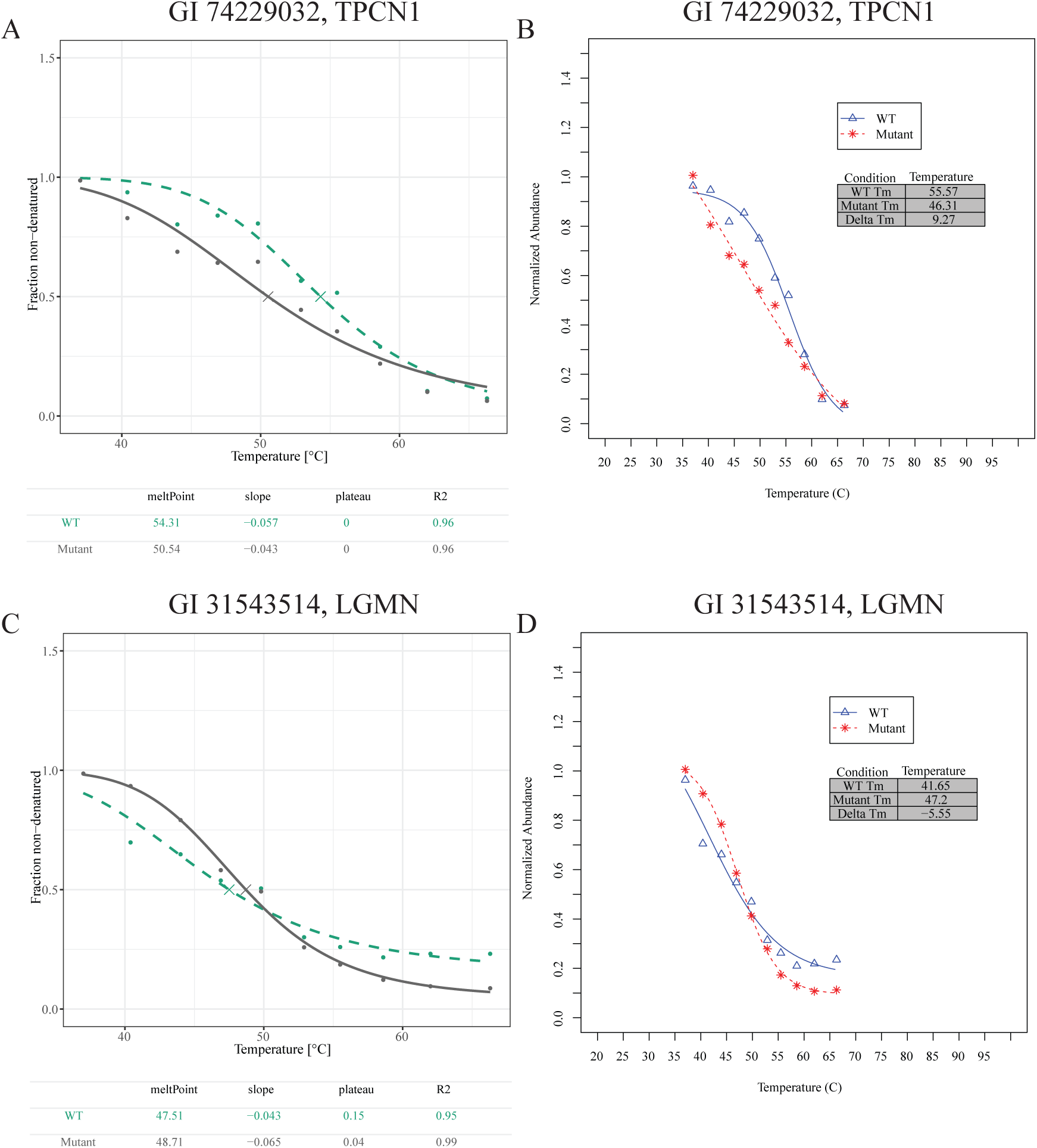
Example rat kidney vs. liver melt shifts from Perrin data sets that were reported significant in our workflow but not significant in the TPP-TR workflow. Melt curves generated from the Two pore calcium channel protein 1, TPCN1 using (A) TPP-TR and (B) our workflow. Melt curves generated from Legumain, LGMN using (C) TPP-TR and (D) our workflow.

### Assessment of step impact on analysis

The melt shift analysis pipeline involves a series of steps that are used to prepare the raw abundance data for analysis, describe the prepared data using fitting routines, and calculate melt temperature shifts from each protein. In order to ascertain the relative influence of each step on the output of the analysis pipeline, a program was written in R which allowed for a multivariate analysis to be executed using various combinations of each step to be run in series with the goal of quantifying the respective output. All 12 data sets described in this article were used in the evaluation. The results from the analysis of the conditions were analyzed using the Fit Model routine in JMP^®^. Fit Model (using a factorial to 2^nd^ degree) was used to describe the observed variability in the outputs as a function of the five factors (Step 1: Exclusion, Step 3: Total Quantitation, Step 4: Curve Fit, Step 6: Curve Fit, Step 8: Melt Definition) used in the study and the relative impact of each pipeline factor on the two outputs was quantified by comparing the scaled estimates of each factor in the model. The fit of each of the four models were good and the order of the effects in importance to the models are summarized in Table 5.

**Table 5.** Summary of factors in order of their relative importance in describing the variability in number of significant proteins and standard deviation of melt temperatures

| Relative Importance | Impacting Total Number of Significant Proteins | Impacting Standard Deviation of Observed Melt Temperature |
| --- | --- | --- |
| 1 (High) | Step 6 Protein Curve Fit | Step 6 Protein Curve Fit |
| 2 | Step 4 Protein Curve Fit | Step 4 Protein Curve Fit |
| 3 | Step 3 Total Quantitation | Step 8 Melt Definition |
| 4 (Low) | Step 8 Melt Definition | Step 3 Total Quantitation |

Results from our optimized workflow show that along with the standard deviation of the melt shift, the second curve fitting routine in step 6 is most important of the variables studied in affecting the number of observed proteins along with the standard deviation of the melt shift. The initial curve fit equation had the next level of importance in the results from our experiment. Interestingly, the use of the 3PL fit for the initial curve fit of the meltome was more beneficial for detecting proteins and reducing standard deviation. This finding of step specific benefit for different curve fitting routines indicates a need for better understanding of how curve fitting equations affect the Tm and melt shift calculations. The definition of the melt temperature and total quantitation had lesser impact on these parameters but were still statistically significant. It should be noted that while the statistics indicate that there is a slight benefit to using the 0.5 definition over the inflection point in the number of proteins observed, we determined that the use of the inflection point over the 50% value allows for analysis of proteins that have non-traditional melt curves where the lower plateau is not equal to 0. The exclusion step used in our *in-silico* experiments did not have a statistically significant impact on the number of proteins or the standard deviation of the melt shifts.

## CONCLUSIONS

Our group investigated the TPP melt shift analysis workflow and the evaluation found that it is beneficial to use the 4PL curve fit over the 3PL fit in order to find proteins with significant melt shifts. To facilitate comparison of our workflow with other data processing pipelines for TPP / CETSA, we have developed the R-based program Inflect. We show also that the number of equation parameters used in curve fitting can dramatically affect the number of proteins observed in the melt shift analysis.

While our work provides extensive insight into the data analysis from TPP experiments, there is still ample opportunity for improvement. One of the challenges of TPP data analysis that needs to be addressed is the explanation of error using measurements of fit that are more suitable for non-linear systems. As more TPP experiments are executed, the experimental procedure will improve along with the methodology for isobaric labeling. As improvements occur, the data analysis pipeline should also be evaluated to determine whether the steps used are most appropriate and beneficial for maximizing the number of biologically relevant results.

## AUTHOR INFORMATION

### Author Contributions

The manuscript was written through contributions of all authors. All authors have given approval to the final version of the manuscript.

### Funding Sources

A portion of the funding for this project was provided by the Indiana University Diabetes and Obesity Research Training Program, DeVault Fellowship (to NAM) by the Showalter Research Trust (to ALM), and by the Indiana University Melvin and Bren Simon Cancer Center Support Grant in support of ALM and ABW (P30CA082709). Additionally, this project was supported in part with support from the Indiana Clinical and Translational Sciences Institute which is funded by Award Number UL1TR002529 from the National Institutes of Health, National Center for Advancing Translational Sciences, Clinical and Translational Sciences Award. The content is solely the responsibility of the authors and does not necessarily represent the official views of funders.

#### ACKNOWLEDGMENT

The authors would like to thank Dr. Ronald Wek for his help in review and discussion of the manuscript. They would also like to thank the other members of the Mosley and Wek labs and the IUSM Proteomics Core for helpful discussions.

#### ABBREVIATIONS

CETSA: Cellular Thermal Shift Assay;
TPP: Thermal Proteome Profiling;
TR: temperature range

